# Resolution Tradeoffs in Modularity Clustering with Application to Single Cell RNA-seq

**DOI:** 10.1101/2025.05.20.655159

**Authors:** Sivan Leviyang

## Abstract

Modularity based clustering was introduced in the network literature for community detection and is now commonly applied to single cell RNA-seq (scRNAseq) datasets for cell type identification. Modularity clustering depends on a resolution parameter, which implicitly determines the number of clusters inferred, but no theory exists describing clustering as a function of the resolution. For scRNAseq, an improperly chosen resolution parameter can lead to erroneous or missed cell types.

In this work, we provide an explicit description of clustering as a function of the resolution parameter through the notion of a splitting resolution, the minimum resolution at which a graph or subgraph is split into multiple clusters. Ve show that the splitting resolution of a subgraph is inversely proportional to the frequency of the subgraph within the graph. This result extends the resolution limit result of Fortunato and Barthelemy to the setting of a general resolution parameter value.

In the network literature, the starting point for modularity clustering is a graph, but in scRNAseq applications the starting point is a cell embedding used to form a graph. Ve show that cell embeddings in scRNAseq can be approximated by Gaussian mixtures. Ve then study splitting resolutions of k-nearest neighbor graphs formed from cell embeddings distributed as a normal or a pair of isotropic normals. For such graphs, we derive formulas for the splitting resolution as a function of sample size, embedding dimension, and the covariance structure of the normals. Ve use our results to provide specific examples of type I vs II error tradeoffs implicit in the choice of the resolution parameter.

---

Unsupervised clustering is commonly applied to single cell RNA-seq (scRNAseq) datasets to identify cell types and states [13]. A scRNAseq dataset is composed of a count matrix, with a row of the count matrix corresponding to a particular cell sampled during the experiment and the columns corresponding to genes. Each entry of the count matrix gives the expression level of a particular gene in a particular cell. Since clustering in a high dimensional space is often ineffective, a dimension reduction such as principal component analysis (PCA) is usually used to embed the rows, or equivalently the cells, of the count matrix in a low dimensional Euclidean space and clustering is applied to the cells in this lower dimensional space.

The particular clustering algorithm applied varies. K-means, hierarchical and mixture model based clustering have been considered [14, 18, 10], but perhaps the most common approach, introduced in [16] and implemented in the popular software packages scanpy [33] and Seurat [27], is to construct a cell graph and then apply modularity clustering. In a cell graph, each node corresponds to a cell and edges connect cells that are near each other in the low dimensional, embedding space. A k-nearest neighbor graph is the simplest way to construct the cell graph, but other algorithms are often used, e.g. a shared nearest neighbor graph [34], UMAP graph [21]. Once the graph has been constructed, the nodes, i.e. cells, of the graph are clustered by maximizing modularity, a clustering metric introduced in the network literature [23]. Given a particular clustering of the graph, modularity is relatively high or low if the number of edges within a cluster is relatively higher or lower than would be expected by chance.

Modularity is parameterized by a resolution parameter, a non-negative scalar that serves to modulate the number of clusters. A high or low resolution parameter leads to relatively many or few clusters, respectively. Within the context of scRNAseq, the resolution parameter implicitly selects a tradeoff in type I/II errors for cell type inference. If we view the alternative hypothesis as the existence of a cell type, then type I and II errors correspond to inferring a cell type which does not exist and failing to infer a cell type that does exist. If the resolution is too high then the clusters may reflect noise rather than capturing true cell types, corresponding to type I errors. On the other hand, if the resolution is too low then individual clusters may combine several cell types, corresponding to a type II error. But while the resolution parameter plays a key role in scRNAseq clustering, no theory exists to guide its selection.

The resolution parameter has also received attention in the network literature. Reichardt and Bornholdt introduced the resolution parameter in analogy to the coupling strength used in spin glass models [26]. Newman connected modularity optimization to maximum likelihood estimation in the context of the stochastic block model (SBM), a probabilistic graph model [24]. In that setting, Newman showed that the resolution parameter can be viewed as selecting a particular parameterization of the SBM. In a different direction, Fortunato and Barthelemy showed that modularity clustering has a resolution limit in the sense that no clusters below a certain size can be identified [7]. However, Fortunato and Barthelemy’s analysis was for a fixed resolution parameter equal to 1 and the implications of their result for settings in which the resolution parameter can be varied is unclear.

In the network literature, modularity is analyzed assuming a particular graph or a graph drawn form a probability distribution over graphs, e.g. an Erdos Renyi graph or a stochastic block model. Modularity clustering of scRNAseq is different because the graph is determined by the cell embeddings. Accordingly, analysis of scRNAseq clustering requires assumption of a particular embedding or a probability distribution of the cell embeddings. In the random geometric graph literature, previous work has mostly focused on spectral clustering of graphs constructed from points sampled from a distribution on a Euclidean space [17, 8]. Modularity clustering has been considered, but in the context of a fixed resolution parameter [3].

In this work, we consider the following question. Given a graph, what is the minimum resolution at which the graph is split into multiple (more than one) clusters! Similarly, given a clustering of a graph at a baseline resolution value, how much higher does the resolution have to be to further split the clusters! We call these minimal resolutions *splitting resolutions* and we derive three novel results characterizing splitting resolutions. For a graph that has been split into several clusters at some low baseline resolution, we show that the splitting resolution for each of these clusters is roughly inversely proportional to the frequency of the cluster. Consequently, low frequency clusters will be split only at high resolution values. This result reframes Fortunato and Barthelemy’s resolution limit in the context of a varying resolution parameter.

We also derive an approximation for the splitting resolution of a knn graph formed from points sampled from a multivariate normal and from points sampled from a mixture of two isotropic normals with different means. Our approximations provide a formula for the splitting resolution in terms of the sample size, embedding dimension, and covariance of the underlying normals. We show that normally distributed samples are relevant to scRNAseq by fitting Gaussian mixture models (GMMs) to the PCA based cell embeddings of scRNAseq datasets and showing that the true splitting resolutions of the datasets are roughly matched by splitting resolutions of simulated datasets generated from the GMMs. We use our approximations to describe a parameter regime in which either a type I or II error is inevitable for modularity based clustering.

Put together, our results provide a quantitative description, in general and on scRNAseq datasets in particular, of the role of the resolution parameter in modulating type I and II errors in modularity based clustering.

## 1 Results

As a starting point, we want to characterize the role of the resolution parameter in selecting a particular clustering. Let *A* be the adjacency matrix for a graph with *n* nodes. Clustering is performed on *A* by maximizing the modularity over all possible clusterings of the graph nodes {1, 2, 3, … , *n*}. Letting 𝒞 represent a generic clustering formed from *L* clusters given by 𝒞_1_, *C*_2_, … , *C*_*L*_, where 𝒞_*ℓ*_ is the set of nodes in the *ℓ*th cluster, modularity is defined as

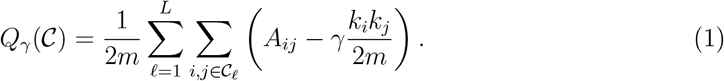

Above, *k*_*i*_ is the degree of node *i*, 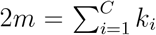 and *γ* is the resolution parameter. For an unweighted knn graph, *k*_*i*_ = *k* and 2*m* = *nk* and heretofore we will make this substitution. The definition of modularity by (1) assumes a configuration model as a null, an assumption we will make throughout.

Rather than maximizing *Q*_*γ*_(𝒞), we can minimize 1 −*Q*_*γ*_(𝒞). A simple algebraic manipulation gives,

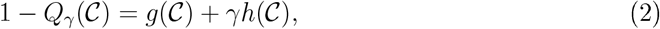

where *g*(𝒞) is the fraction of edges that start in one cluster of 𝒞 and end in another and 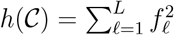 where *f*_*ℓ*_ is the fraction of nodes in cluster *ℓ*. (These definitions for *g*(𝒞) and *h*(𝒞) assume an unweighted knn graph. If *A* is the adjacency matrix of a generic weighted graph, then the decomposition (2) still holds with *g*(𝒞) and *f*_*ℓ*_ expressed in terms of edge weights.) The decomposition (2) frames modularity clustering as a graph cut problem. The standard graph cut problem seeks to split a graph into two clusters that minimize the ratio of edges that cross the clusters to the size of the clusters, see [32, 22] for examples of the graph cut problem in the context of geometric random graphs. Typically, the graph cut problem is considered with *γ* = 1 in the notation of (2).

The minimization,

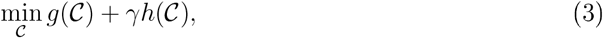

frames modularity maximization in terms of the minimization of two objectives *g* and *h* and we can interpret *γ* as selecting a point on a Pareto frontier. More precisely, since there are only finitely many clusterings, let (*g*_*i*_, *h*_*i*_) for *i* = 0, 1, 2, 3, … , *m* be the *g*(𝒞), *h*(𝒞) values for some clustering 𝒞 that has maximal modularity for some *γ* ∈ (0, ∞), where *m* is the total number of optimal clusterings. The (*g*_*i*_, *h*_*i*_) form a Pareto frontier, meaning that no clustering 𝒞 can achieve *g*(𝒞) *< g*_*i*_ and *h*(𝒞) *< h*_*i*_ since then (*g*_*i*_, *h*_*i*_) would not form an optimal solution. It’ll be useful to assume that the *g*_*i*_ are in increasing order, which puts the *h*_*i*_ in decreasing order. Figure 1 shows the modularity frontier of a scRNAseq dataset.

**Figure 1.**
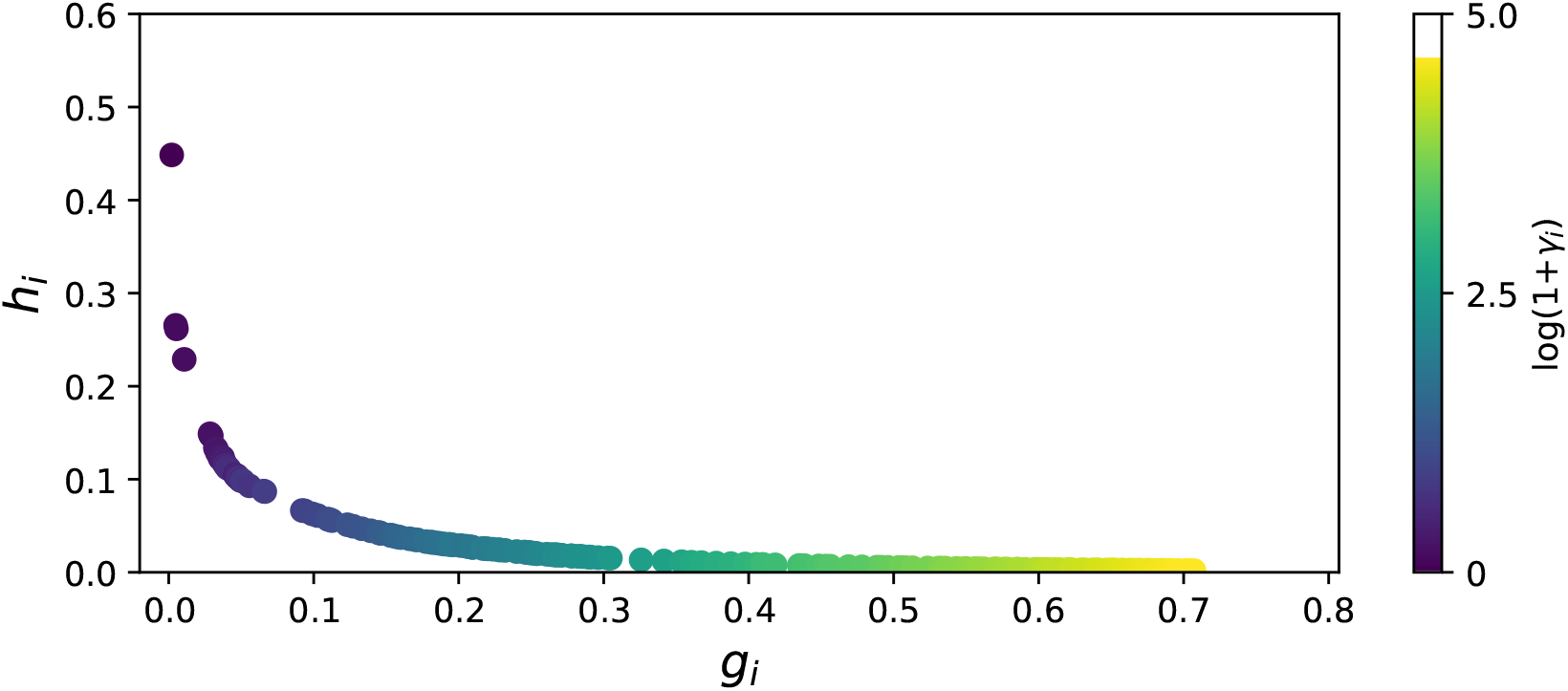
Pareto Modularity Frontier for a scRNAseq Dataset. Shown are (*g*_*i*_, *h*_*i*_) and *γ*_*i*_ that form the modularity frontier for a scRNAseq dataset sampled from heart tissue [12]. Colors give *γ*_*i*_. Each (*g*_*i*_, *h*_*i*_) corresponds to an optimal clustering of the cells at resolution *γ*_*i*_. For this optimal clustering, *g*_*i*_ is the fraction of edges that cross clusters and *h*_*i*_ is the squared sum of the cluster frequencies.

For sufficiently small *γ*, the optimal clustering is formed by the disconnected components of the graph, which gives *g*_0_ = 0. If the graph is connected, *h*_0_ = 1 (since all nodes are in one cluster making *L* = 1 and *f*_1_ = 1), otherwise *h*_0_ *<* 1. For Figure 1, *g*_0_ = 0 and *h*_0_ ≈ 0.45. For sufficiently large *γ*, the optimal clustering places a single node in each of *n* clusters giving *g*_*m*_ = 1 and *h*_*m*_ = 1*/n*.

In the case *i* ≠ 0, for each (*g*_*i*_, *h*_*i*_) there exists a *γ*_*i*_ such that min_*C*_ 1 − *Q*_*γ*_(𝒞) = *g*_*i*_ + *γh*_*i*_ for *γ* ∈ [*γ*_*i*_, *γ*_*i*+1_]. The *γ*_*i*_ form an increasing sequence. At the resolution *γ*_*i*_, *g*_*i*−1_ + *γ*_*i*_*h*_*i*−1_ = *g*_*i*_ + *γ*_*i*_*h*_*i*_, which characterizes *γ*_*i*_ for *i >* 0 as a function of the (*g*_*i*_, *h*_*i*_),

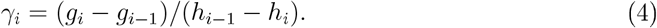

See the Methods for a proof of the above claims.

Modularity is typically framed as the difference of the observed number of internal edges in a cluster to the expected number of internal edges, which are given by the *A*_*ij*_ and *k*_*i*_*k*_*j*_*/*2*m* terms in (1), respectively. It is however unclear how to incorporate the resolution parameter into this framework. Reichardt and Bornholdt defined the resolution parameter as parameterizing the Hamiltonian of a spin glass model [26]. Viewing modularity optimization as a penalized optimization provides an alternative description for the role of the resolution parameter which allows for an explicit connection between the resolution parameter and clustering properties through (4).

### 1.1 Splitting Resolutions for Baseline Clusters

In [7], Fortunato and Barthelemy note that modularity clustering typically forms clusters which would be further clustered if considered as separate graphs. Fortunato and Barthelemy show that this problem reflects a failure of modularity clustering to operate across scales. In particular, Fortunato and Barthelemy show that modularity clustering cannot identify clusters with less than 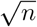 nodes (unless the clusters are completely disconnected from the other nodes in the graph). The results of Fortunato and Barthelemy assume *γ* = 1.

We now derive a similar result but in the case of a general *γ*. Suppose that we cluster a graph at a relatively low *γ* (e.g. below we’ll choose *γ* = 0.1). We refer to the clusters formed at this *γ* as the baseline clusters. Our goal is to determine the *γ* at which a baseline cluster will be subclustered, i.e. split into multiple clusters.

To put this in the context of scRNAseq, we apply modularity clustering to seven scRNAseq datasets with *γ* = 0.1, which is below the default resolutions of *γ* = 0.8 and *γ* = 1 used in Seurat and scanpy. Four of the datasets are cell atlases from the tabula sapiens consortium [12] for blood, eye, heart, and tongue tissue, respectively. The other three datasets sample PBMC [35], brain [28] and pancreas [6] tissues, respectively. See the Supporting Information for further details. Shown in Figure 2 are the baseline clusters for the three of the datasets. The clusters generated at *γ* = 0.1 are roughly disconnected with less than 1% of edges crossing clusters. The baseline clustering roughly corresponds to cell types, but clusters sometimes contain multiple cell types and single cell type are sometimes split into multiple clusters.

**Figure 2.**
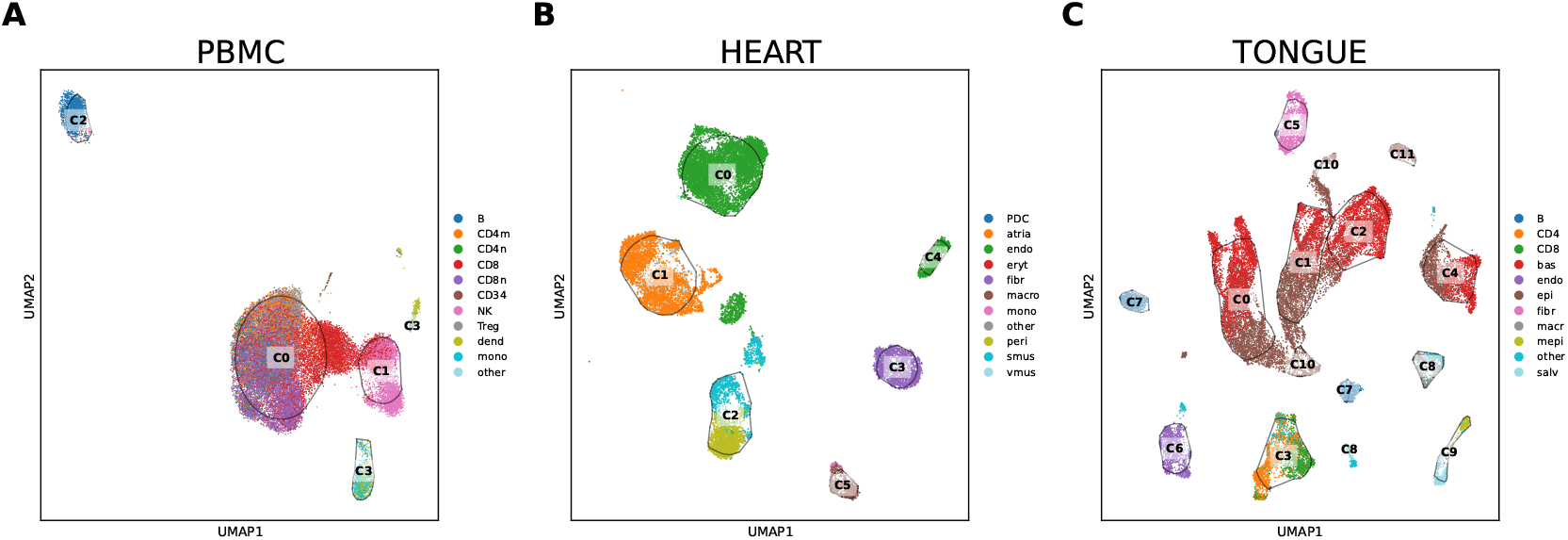
Baseline Clustering UMAP. visualization of modularity clustering at *γ* = 0.1 for three scRNAseq datasets (PBMC [35] and heart and tongue atlases [12]). Clusters are delineated by black boundaries and text labels. Individual cells (i.e. the scatter points) are colored by their cell types.

Returning to the generic case, for each baseline cluster define the cluster’s global splitting resolution *γ*_*s*_ as the minimum resolution at which the baseline cluster is split into multiple clusters. Importantly, although *γ*_*s*_ is determined by splitting of a particular baseline cluster, we are still clustering on the full graph. The existence of a finite Pareto modularity frontier guarantees the existence of *γ*_*s*_. Each baseline cluster has its own *γ*_*s*_ and we compute *γ*_*s*_ numerically by running the Leiden [31] algorithm on the adjacency matrix *A* on a grid of *γ* values. Although *γ*_*s*_ applies to a particular baseline cluster, *γ*_*s*_ depends on the full adjacency matrix *A*, which motivates the term global splitting resolution.

Next, for each baseline cluster define a local splitting resolution, 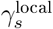. Define the local adjacency matrix *A*^local^ as the adjacency matrix of the local graph that would be constructed if the dataset was restricted to nodes in the baseline cluster. Then 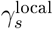 is defined as the minimal resolution at which the baseline cluster splits into multiple clusters using *A*^local^. We can compute 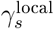 numerically by constructing the local graph and running a Leiden algorithm at different resolutions. Note that each baseline cluster has its own pair of global and local splitting resolutions: *γ*_*s*_ and 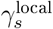.

If a baseline cluster forms a disconnected component relative to the rest of the cell graph, then the global and local splitting resolutions are related by,

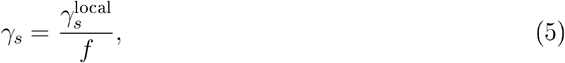

where *f* is the fraction of cells in the dataset that are in the baseline cluster. (Recall, we assume the graph is a knn graph. For a general weighted graph, 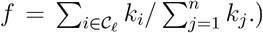 To understand the intuition behind (5), let *n* _*b*_ be the number of nodes in a baseline cluster. Roughly, the cluster’s local and global splitting resolutions are determined by modularity of the form (1) but with 2*m* = *n*_*b*_*k* and 2*m* = *nk*, respectively, leading to the relation 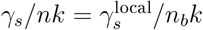 which gives (5). A complete proof of (5) is given in the Methods.

Above we defined the splitting resolution as the minimum resolution for which a baseline cluster is split, but the split may involve a small number of nodes - for example the baseline cluster could be split into two clusters composing 99% and 1% of the nodes. To address such an effect, we can redefine the splitting resolution as the minimal resolution at which clustering achieves a certain rand index relative to the trivial clustering of a single cluster containing all nodes. In this case, (5) still holds using identical arguments.

(5) becomes an approximation rather than an equality if the baseline clusters are roughly disconnected. We assess the accuracy of this approximation using scRNAseq data. Figure 3 shows the estimated and true values for *γ*_*s*_ over the seven scRNAseq datasets. There are 55 baseline clusters in total and for each we compute the true *γ*_*s*_ and *γ*_*s*_ as predicted by (5) given the frequency of the baseline cluster *f* and 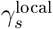. As can be seen in the figure, (5) provides a good approximation of *γ*_*s*_.

**Figure 3.**
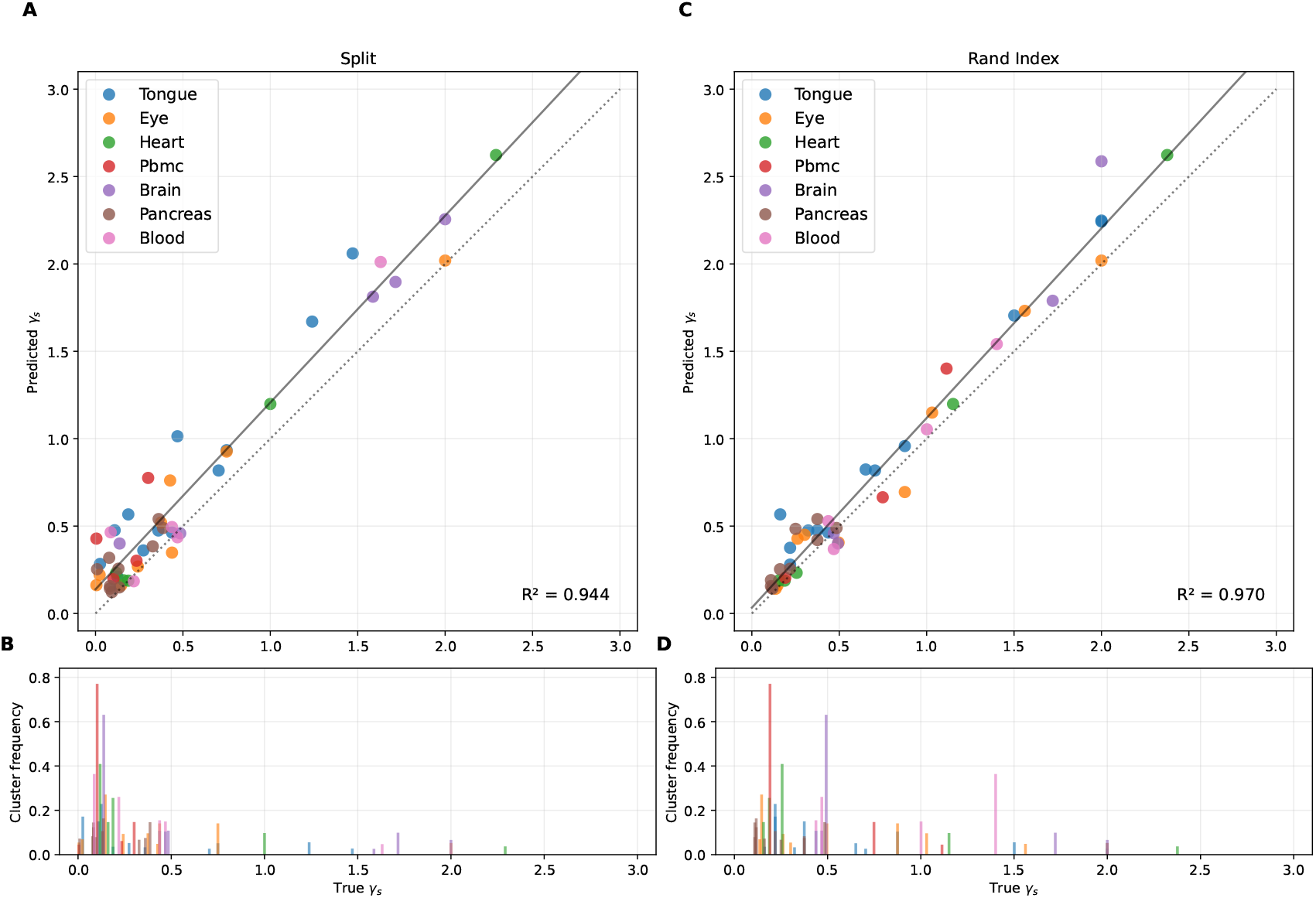
Prediction of Global Splitting Resolution. (A,C) Each point shows the predicted global splitting resolution based on (5) vs the true global splitting resolution for a single baseline cluster within one of the 7 datasets. The predicted splitting resolution is computed using (5). (B,D) The height of each bar shows the frequency of a baseline cluster. The x-value of the bar is the true global splitting resolution and corresponds to the x-value of a point in the panel above. In the left panels and right panels, splitting resolution is defined as the minimum resolution that splits the baseline cluster in any way and with a rand index below 0.8, respectively. The dashed and solid black lines in the upper panels are the 45 degree line and the regression line for the scatter points, respectively.

The Fortunato and Barthelemy result can be recast in terms of the resolution parameter and (5). A baseline cluster will be split at a resolution that is inversely proportional to the fraction of graph nodes in the baseline cluster which leads to a problem of scales. If we want to cluster small components, we need *γ* to be large. However, as we show below, a large *γ* can lead to clustering of graph components that do not have any clustered structure.

(5) has significant implications in the context of scRNAseq. Consider baseline cluster 2 of the PBMC dataset which is composed mostly of B cells. The local splitting resolution of the cluster is 0.38. If our dataset was formed just of B cells, this cluster would be split at the default resolution of Seurat, *γ* = 0.8. However, the cluster forms 5.5% of the full dataset, giving a global splitting resolution of roughly 7; a resolution that is far beyond the default resolution.

At a given resolution, a cell type may be split into multiple clusters, which can be viewed as a type I error, and multiple cell types may be clustered together, which can be viewed as a type II error. To prevent the splitting of a cell type into multiple clusters, resolution has to be lowered below the global splitting resolution. To force the splitting of cell types that are clustered together, resolution has to be raised above the global splitting resolution. Clearly, situations can arise in which we can’t eliminate both error types, especially since the global splitting resolutions depend on the frequency of the cell types.

### 1.2 Splitting Resolutions for Normal Embeddings

(5) describes global splitting resolutions as a function of local splitting resolutions. To determine local splitting resolutions, we need a statistical description of the knn graph. In this context,recall that for scRNAseq datasets the knn graph is formed from samples *x*_*i*_ in the PCA embedding space. We assume the *x*_*i*_ are drawn independently from a probability distribution on ℝ ^*p*^, which implicitly defines a probability distribution on the resulting knn graph. For these embeddings, we define the splitting resolution as the minimum resolution at which the full graph is split into more than one cluster. In terms of our previous terminology, this is the global splitting resolution based on a cluster that is the whole graph, but below we simply call this resolution the splitting resolution.

We will consider two particular distributions for the *x*_*i*_. First, the *x*_*i*_ are drawn indepen-dently from a multivariate normal with mean *µ* and covariance Σ, i.e. *x*_*i*_ ∼ 𝒩 (*µ*, Σ). Second, the *x*_*i*_ are drawn from a mixture model composed of two isotropic multivariate normals with different means,

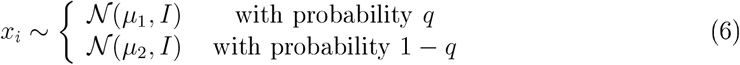

where *µ*_1_, *µ*_2_ ∈ *ℝ* ^*p*^, *I* is the *p* × *p* identity matrix, and *q* is the probability of drawing from the first mixture.

To motivate the choice of sampling *x*_*i*_ from normal distributions, we fit Gaussian (i.e. normal) mixture models (GMM) to the *x*_*i*_ forming the baseline clusters in the seven scRNAseq datasets. For each baseline cluster, we used the Python sklearn package to fit a GMM to the baseline cluster’s *x*_*i*_ with the number of mixtures varying from 1 to 10. We then used the Akaike information criterion (AIC) to to select the number of mixtures. To verify that a GMM is a reasonable model for analyzing local splitting resolutions, we compared the true local splitting resolution of the baseline clusters to a simulated local splitting resolution based on the best fit GMM model. The simulated splitting resolution was generated by sampling *x*_*i*_ from the fitted GMM, building a knn graph, and computing *γ*_*s*_. Figure 4 shows that the GMM models are reasonable predictors of the splitting resolutions.

**Figure 4.**
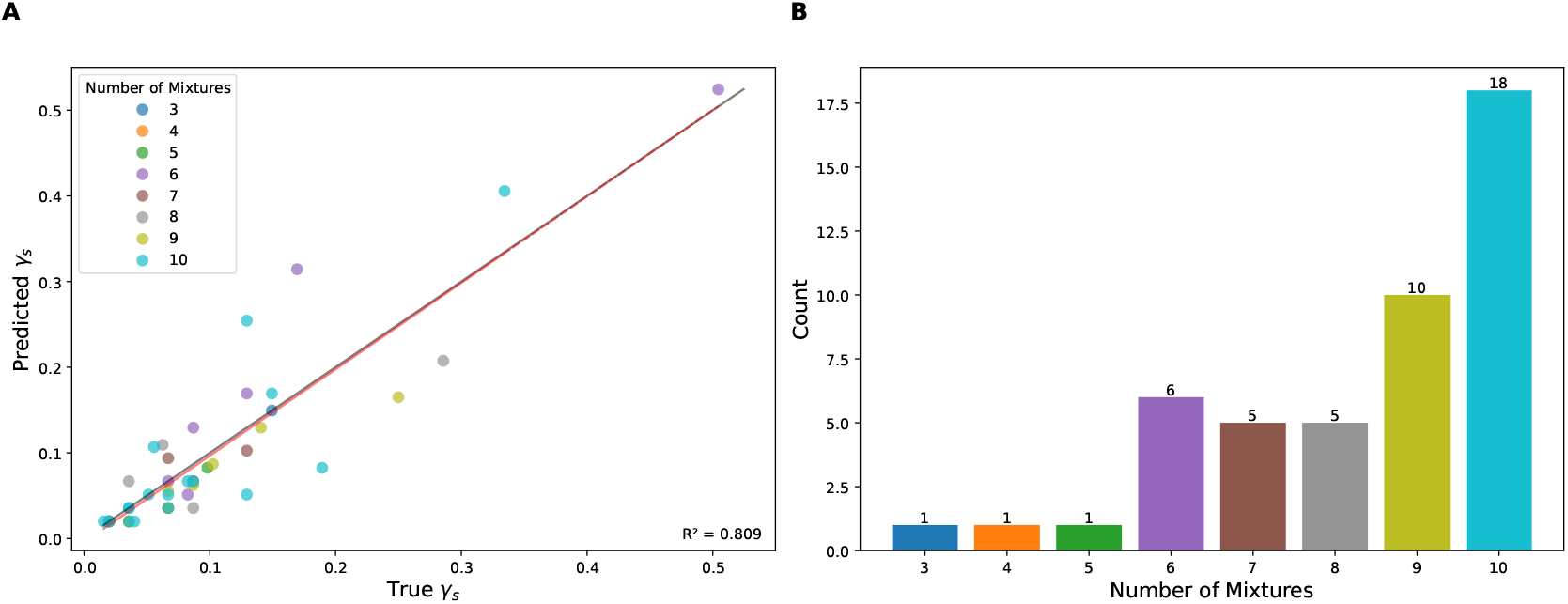
Local Splitting Resolutions of GMMs. (A) The splitting resolution generated by the best fit GMM vs the true splitting resolution. Each point corresponds to a particular baseline cluster within a dataset. For each baseline cluster, GMM mixtures with 1 *−* 10 mixtures were fit to the data and the best fit GMM was determined using the AIC. (B) The number of mixtures composing the best fit GMMs. For example, 18 of the best fit GMMs have 10 mixtures.

### 1.3 Splitting Resolution for a Normal

Let *x*_*i*_ ∼ 𝒩 (*µ*, Σ) for *i* = 1, 2, … , *n* with each *x*_*i*_ ∈ *R*^*p*^. For a knn graph, edges are determined by the distances between the samples *x*_*i*_. We can therefore take *µ* = 0. Let *X* be the row binding of the *x*_*i*_ samples, making *X* an *n* × *p* matrix.

Suppose we construct a knn graph from *X*. We will assume that the graph cannot be partitioned into disconnected components. In general, this need not be the case if the samples have large variance or *n* is small, but we have found this assumption to hold over the parameter spaces we simulate below. Further work is needed to clarify the exact parameter regime in which the assumption holds.

Given this assumption, suppose we partition the nodes of the knn graph into two clusters. For any particular sample *x*_*i*_, let *e*_*i*_ be the fraction of *x*_*i*_’s neighbors that do not belong to *x*_*i*_’s cluster. If we assume the clustering is optimal, in the sense of being on the Pareto modularity frontier, then the splitting resolution satisfies *γ*_*s*_ = *γ*_1_ and (4) gives,

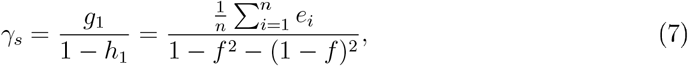

where *f* and 1 − *f* are the fraction of samples in each of the two clusters. A law of large numbers argument then suggests the approximation,

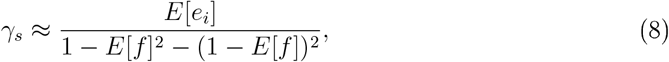

and approximating the splitting resolution reduces to estimating *E*[*e*_*i*_] and *E*[*f* ]. Using this approach, in the Methods we derive the following approximation for the splitting resolution *γ*_*s*_.

#### Approximation 1

*Let E*_*1*_, *E*_*2*_, *E*_*3*_, … *be independent exponential random variables where E*_*i*_ *has mean 1/i. Let W*_*1*_, *W*_*2*_, *W*_*3*_, … *be iid standard Normals and et W be an independent standard Normal* . *Based on these sequences of random variables de ne the sequence D*_*j*_ *for j = 1, 2, 3*, … *as*,

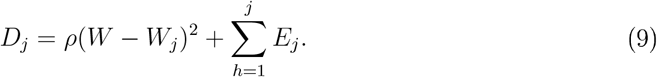

*where*

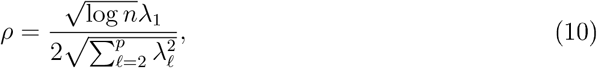

*and where λ*_*1*_, *λ*_*2*_, … , *λ*_*p*_ *are the eigenva ues of Σ. Then for n ≫ p*,

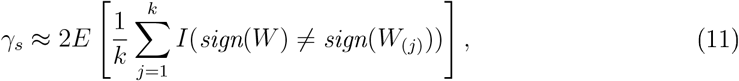

*where (j) are the order statistics of the D*_*j*_, *i*.*e. D*_*(1)*_ *is the D*_*j*_ *with the sma est va ue, D*_*(2)*_ *the next sma est, and so on*.

(In the methods, we also present an approximation that relaxes the condition *n* ≫ *p*.)

Approximation 1 is useful because it expresses *γ*_*s*_ as a function of *ρ* which in turn is expressed in terms of the eigenvalues of Σ, *n* and *p*. We do not have an analytic expression for the right side of (11) as a function of *ρ*, but it can easily be estimated by Monte Carlo.

We assess the accuracy of Approximation 1 in two ways. First, we construct different covariance matrices Σ, generate *n* samples *x*_*i*_ from 𝒩 (0, Σ), calculate the true *γ*_*s*_ by constructing a knn matrix and applying the Leiden algorithm and then compare the true value of *γ*_*s*_ to Approximation 1. We construct Σ as a diagonal matrix with *p*− 1 entries uniformly chosen on [1, 5] and one entry chosen uniformly on [5, 100], leading to different eccentricities and varying *ρ*. (Making Σ diagonal is not a limitation since the knn graph is invariant to rotations and any covariance matrix can be rotated to become diagonal.) We generate 100 such Σ for all nine combinations of *n* = 1000, 5000, 10000 and *p* = 20, 50, 100 giving us 900 comparisons of the approximated *γ*_*s*_ to the true *γ*_*s*_. We also consider the approximation of *γ*_*s*_ based on the covariance matrices generated by the GMM fits of the scRNAseq datasets. In the case of these covariance matrices, of which there are 336, *n* is determined by the baseline cluster size and *p* = 50. Figure 5 shows that Approximation 1 is a good estimator for the true *γ*_*s*_, at least for the covariance matrices we consider.

**Figure 5.**
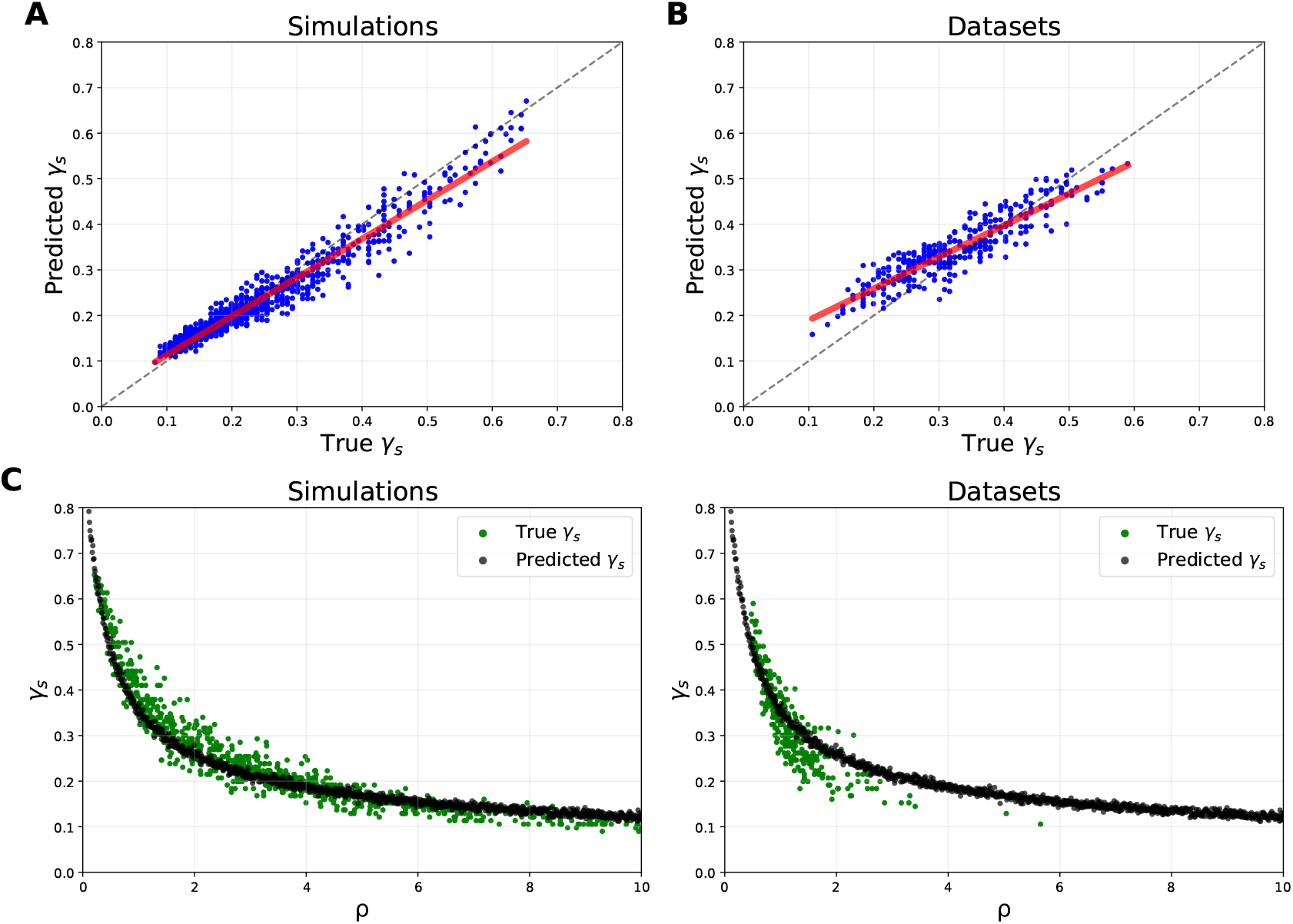
Approximated vs True Splitting Resolution of a Normal. (A,B) The blue points compare the splitting resolution *γ*_*s*_ predicted by Approximation 1 to the true splitting resolution. The red and black lines are the regression line and 45 line respectively. Each point in panel A and B is generated by constructing a covariance matrix Σ and using a Σ from the GMMs, respectively. (C,D) The same data as (A,B) but shown as a function of *ρ*. See main text for details regarding *n, p* values.

To understand Approximation 1 in a simple context, set *k* = 1, making

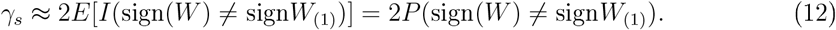

Consider the case of *ρ* = 0 (or *ρ* small). In this case 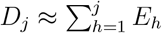 and the *W*_*j*_ and *W* have no impact on the order statistics of the *D*_*j*_, making *W*_(1)_ and *W* independent. Since *W*_(1)_ and *W* are independent, *P* (sign(*W*) ≠ sign(*W*_(1))_) = 1*/*2, making *γ*_*s*_ = 1, meaning that the *x*_*i*_ will not be split into more than one cluster for any *γ <* 1 and will be split for any *γ >* 1.

Now consider the case of *ρ* → ∞ (or *ρ* large). In this case *D*_*j*_ ≈ *ρ*(*W* − *W*_*j*_)^2^ and the order statistics of the *D*_*j*_ are determined by the *W*_*j*_. *W*_(1)_ minimizes the distance to *W* over all *W*_*j*_. In this case *W*_(1)_ ≈ *W* and *P* (sign(*W*) *≠* sign(*W*_(1))_) ≈ 0, making *γ*_*s*_ ≈ 0, meaning that the *x*_*i*_ will be split into more than one cluster at any *γ >* 0.

It’s instructive to consider an isotropic normal for which *λ*_*i*_ = 1 for *i* = 1, 2, … , *p*. In that case, 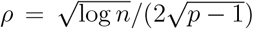. If we associate *p* with the PCA dimension applied to a scRNAseq dataset and *n* with the number of cells and set *p* = 50 and *n* = 10^4^ then *ρ* = .22 which gives *γ*_*s*_≈ 0.66. Clearly, clustering an isotropic normal reflects a type I error, suggesting that a resolution value of 0.66 is too high. Figure 6 visualizes clustering of normals at resolution *γ* just below and just above the splitting resolution *γ*_*s*_. The top panels (A,B) show clustering of an isotropic normal at 0.61 and 0.71, just below and above *γ*_*s*_ = 0.66. As predicted, modularity clustering at *γ* = 0.71 splits the samples into two clusters while clustering at *γ* = 0.61 leads to a single cluster containing all samples.

**Figure 6.**
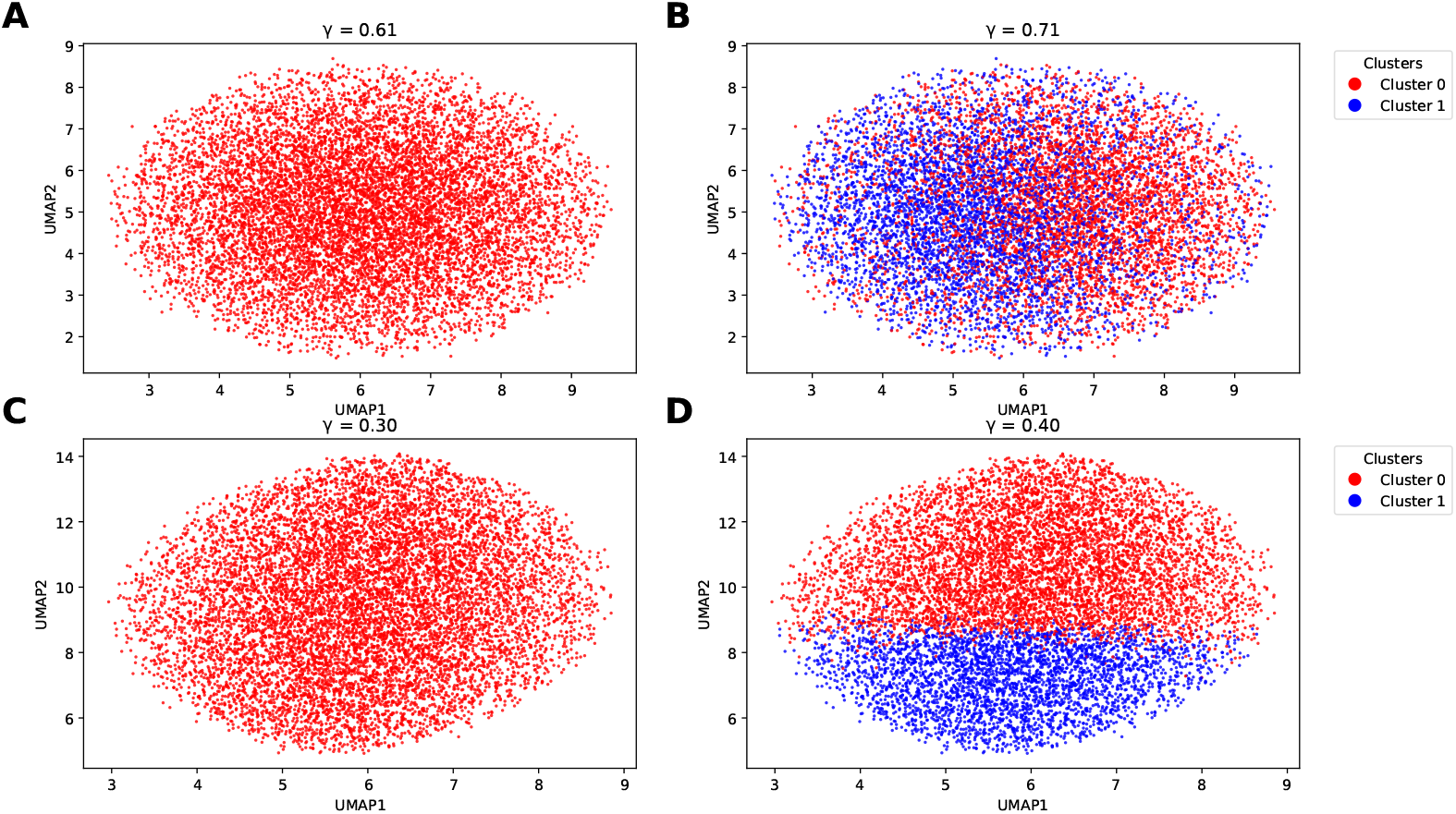
Modularity Clustering of a Normal. Shown are UMAP visualizations of (A,B) an isotropic normal with variance 1 along all coordinates and (C,D) a normal with variance 1 along all but one coordinate for which the variance is 5. Each point represents a sample *x*_*i*_. For both normals *n* = 10000 and *p* = 50. By Approximation 1, *γ*_*s*_ = 0.66 and *γ*_*s*_ = 0.35 for the samples from the isotropic and non-isotropic normals, respectively. When the resolution *γ* is (left panels) 0.05 less and (right panels) 0.05 greater than *γ*_*s*_, modularity clustering infers a single and two clusters, respectively.

### 1.4 The Splitting Resolution of a Pair of Isotropic Normals

For *x*_*i*_ drawn from a pair of isotropic normals, invariance of the knn graph relative to shifts allows us to assume *µ*_1_ = *µ* and *µ*_2_ = −*µ*. Given this parameterization, in the Methods we derive the following approximation for *γ*_*s*_.

#### Approximation 2

*Let E*_*j*_, *W*_*j*_ *and W be as in Approximation 1. Let B*_*j*_ *and B be independent Rademacher random variables with success probability q (recall q is the probability of drawing from mixture 1* . *Define*,

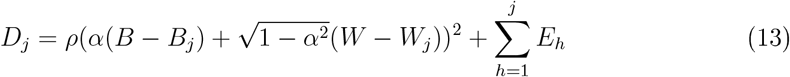

*where*

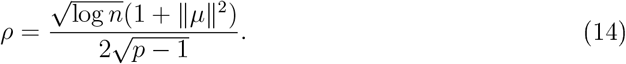

*and*

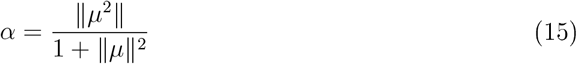

*Then for n ≫ p*

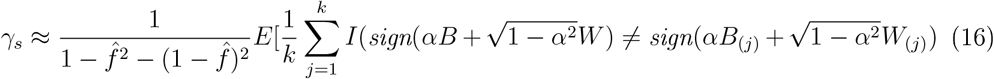

*where (j) are the order statistics of the D*_*j*_ *and*

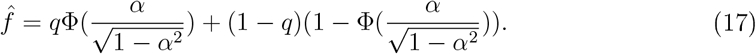

Φ *is the standard Normal cdf*.

Approximation 2 is more complex than Approximation 1. In particular, there are now two parameters *ρ* and *α*; however both are functions of ∥*µ*∥. For orientation, think of *γ*_*s*_ as a function of ∥*µ*∥ with *n, p* and *q* fixed. If ∥*µ*∥ = 0 then *α* = 0, 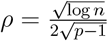 , and 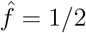 and Approximation 2 is identical to Approximation 1 with Σ = *I*. For ∥*µ*∥ large, *α* = 1, *ρ* is large, and 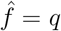, making *D*_*j*_ ≈ *ρ*(*B* − *B*_*j*_)^2^. Similarly to the case of large *ρ* in Approximation 1, *B*_(1)_ ≈ *B* and *γ*_*s*_ ≈ 0.

Figure 7 compares the prediction for *γ*_*s*_ using Approximation 2 to the true *γ*_*s*_ over different values of *µ*. For each of the 27 combinations of *n* = 1000, 5000, 10000; *p* = 20, 50, 100; and *q* = 0.5, 0.75, 0.95 we generated 100 values of ∥*µ*∥ equally spaced between 0 and 4. Approximation 2 is relatively accurate, although there is a clear overestimate for small values of ∥*µ*∥ (large *γ*_*s*_) and there is noise in the estimator. Figure 8 visualizes samples for a pair of isotropic normals with ∥*µ*∥ = 2 at a resolution just above and below *γ*_*s*_.

**Figure 7.**
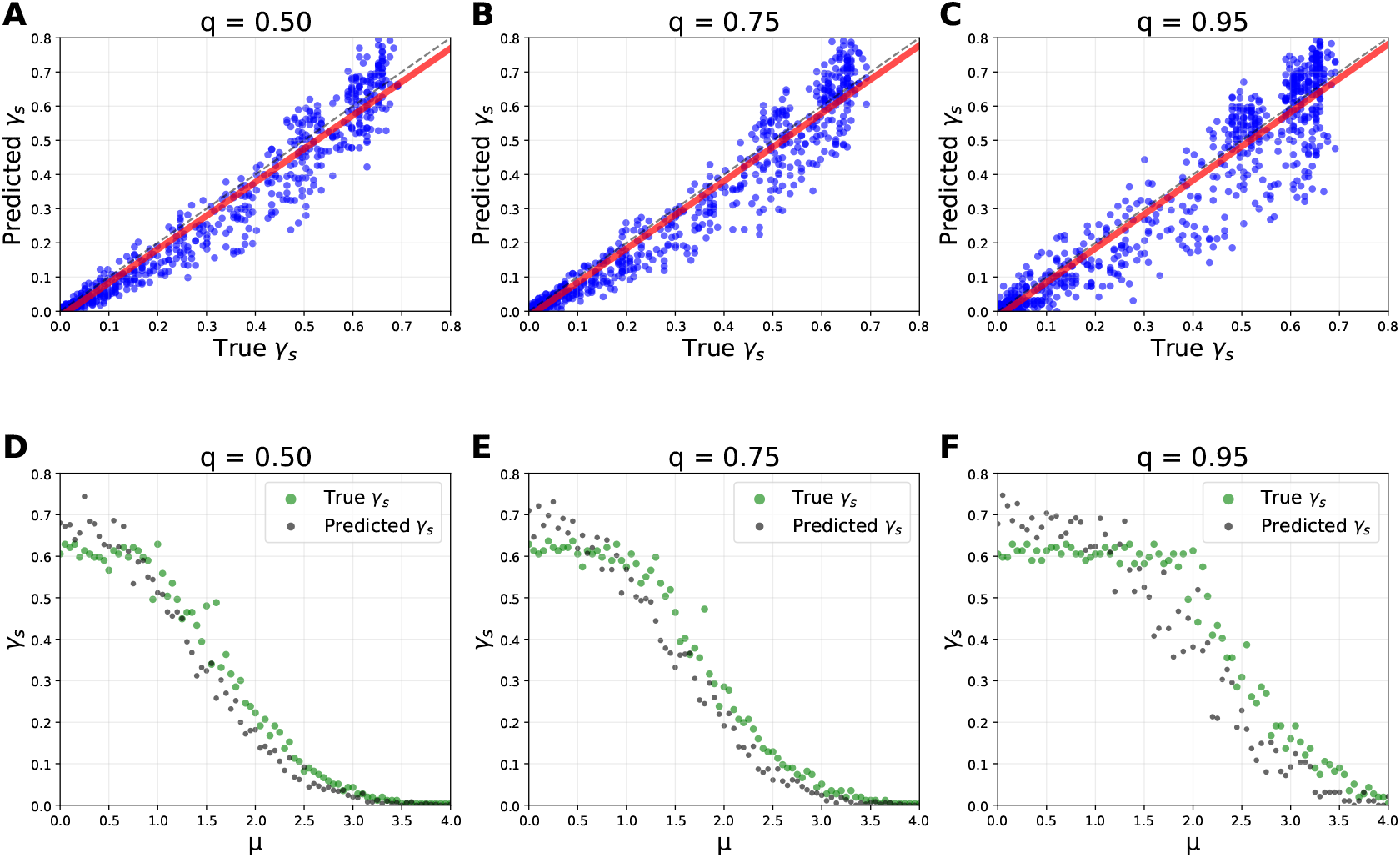
Approximation of the Splitting Resolution for a Pair of Isotropic Normals. (A,B,C) Splitting resolution predicted by Approximation 2 versus the true splitting resolution for the different mixture frequencies *q* over *n* = 1000, 5000, 10000 and *p* = 20, 50, 100. (D, E, F) The same data as (A,B,C) but shown as a function of *µ* and restricted to *n* = 1000 and *p* = 50.

**Figure 8.**
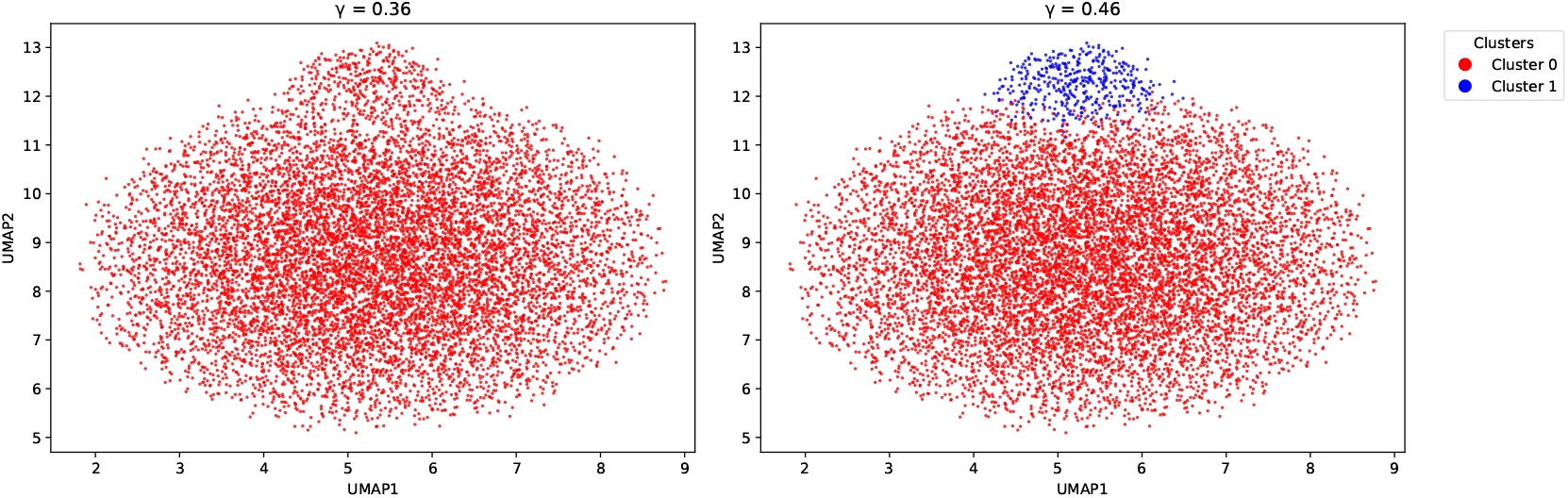
Modularity Clustering of a Pair of Isotropic Normals. Shown is UMAP visualization of a pair isotropic normals parameterized by ∥*µ*∥ = 2, *n* = 10000, *p* = 50 and *q* = 0.95. Each point represents a sample *x*_*i*_. When the resolution *γ* is (A) 0.05 less and (B) 0.05 greater than *γ*_*s*_ = 0.41, modularity clustering infers a single and two clusters, respectively.

Putting the two approximations together allows us to concretely describe an inherent limitation of modularity optimization in terms of type I/II errors. Consider a dataset that merges samples from a single normal with samples from a pair of isotropic normals. For simplicity, assume that the single normal has a diagonal Σ with Σ_*ii*_ = 1 for all *i* ≠ 0 and Σ_11_ = *σ*^2^ *>* 1 and that the number of *x*_*i*_ drawn from each of the two models is equal. If we assume that the mean of the single normal is far from the mean of the isotropic normals, then a knn graph formed from the data will create two disconnected components of equal size, composed of samples from the single normal and pair of normals, respectively.

By (5) 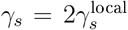 since *f* = 1*/*2 and Approximations 1 and 2 give 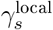. Ideally, we would like to find a resolution *γ* that would not split the single normal, since that would be a Type I error, but that would split the two paired normals, since not doing so would be a Type II error. Figure 9 describes the region of the parameter space formed from *σ*^2^, ∥*µ*∥ that avoids both type I and type II error and the region for which either a type I or a type II error cannot be avoided. At ∥*µ*∥ = 1.5 and *σ*^2^ = 17, the global splitting resolutions of the single normal is 0.44, which is lower than the global splitting resolution of 0.77 of the paired normals, meaning that clustering the pair requires a type II error of splitting the single normal.

**Figure 9.**
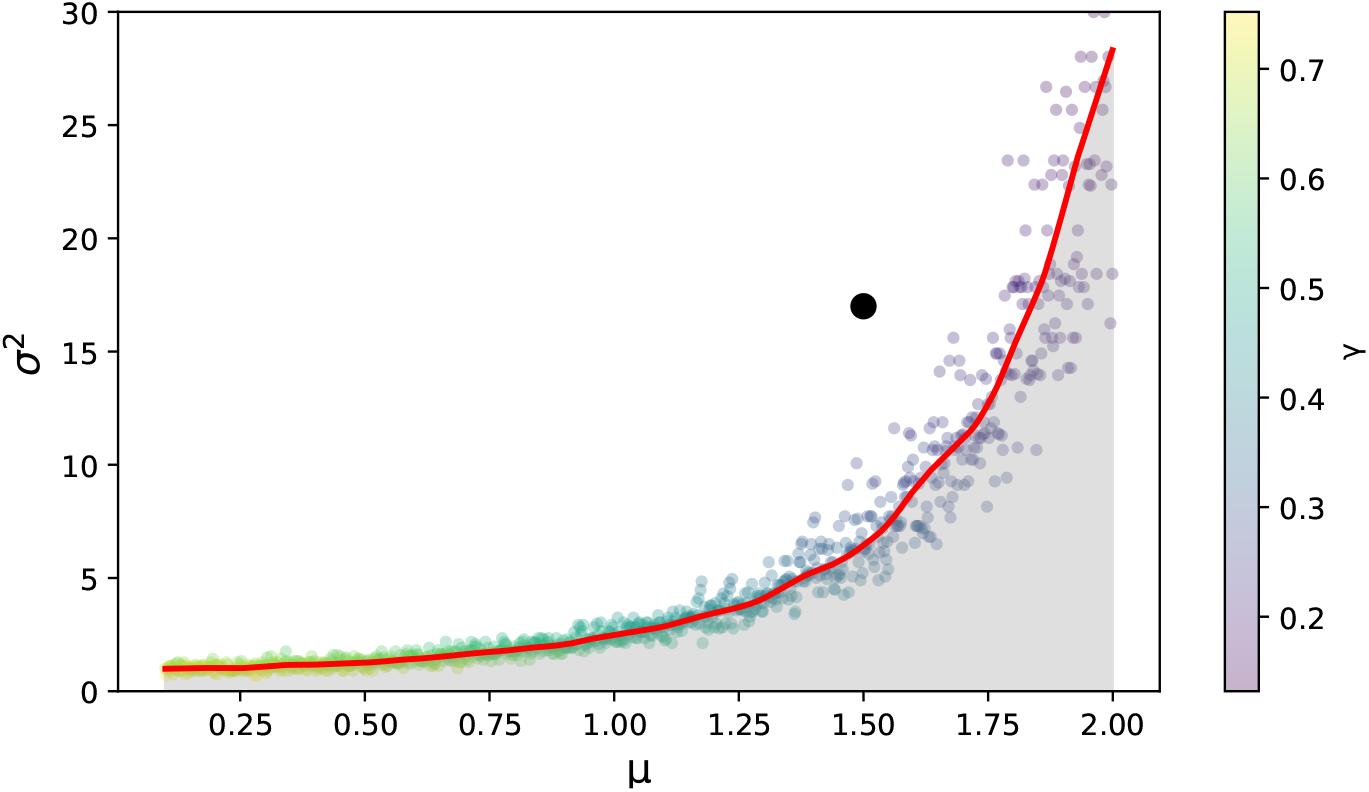
The Resolution Parameter and Type I/II Errors. The points show *σ*^2^, ∥*µ*∥ combinations for which the global splitting resolution of the single normal and two paired isotropic normals are equal. The points are colored by this splitting resolution. *n* = 1000 and *p* = 50 for both models. For the isotropic pair model, *q* = 0.5. For the single normal, Σ is diagonal with all entries 1 except for one entry *σ*^2^. Above the curve formed by these points (the white region), it is impossible to find a resolution *γ* for which the paired normals can be clustered without splitting the single normal into multiple clusters, i.e. either a type I or type II error will be made. Below the curve (the grey region), a resolution exists that clusters the paired normals but not the single normal. The black point specifies the ∥*µ*∥ , *σ*^2^ combination mentioned in the text.

## 2 Discussion

Modularity clustering is a popular approach for identifying cell types in scRNAseq datasets, but little theory exists to guide its application. In particular, clustering has an inherent tradeoff between identifying too few clusters, which do not capture the full structure of the dataset, and identifying too many clusters, which infer structure that does not exist in the dataset. In this work, we provide an analysis that describes how the resolution parameter shapes this tradeoff.

Many scRNAseq datasets are formed of multiple cell types. Cell graphs formed from these type of datasets are composed of roughly disconnected components that can be clustered at a low resolution parameter. We cluster 7 scRNAseq datasets at *γ* = 0.1. In each case, this baseline clustering leads to clusters that roughly connect with cell type. We investigate the resolution value above 0.1 at which a baseline cluster would be split and we show that this resolution varies inversely with the frequency of the baseline cluster.

Fortunato and Barthelemy showed that small roughly disconnected components cannot be clustered at a resolution of 1. If the resolution parameter can be varied, then any size component can be clustered if it is sufficiently disconnected. However, our result shows that the resolution parameter needed must increase as the component’s frequency drops. From the perspective of scRNAseq, our results shows that we can identify rare cell types by raising the resolution, but likely at the expense of over-clustering more common cell types.

To put this observation on a firmer footing, we derive the local splitting resolution for a normal and pair of isotropic normals. For the normal model we show that the splitting resolution is a function of a parameter *ρ* which in turn depends on sample size *n*, embedding dimension *p* and the covariance of the normal. Normals with large eccentricity will have low splitting resolutions which, assuming we don’t want to cluster normals, will lead to type I errors. We find that the dependence of *ρ* on sample size *n* is proportional to 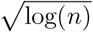, suggesting that increasing sample size will not fundamentally change the splitting resolutions and therefore not eliminate these type I errors. For example, going from 100, 000 cells to 1 billion cells will decrease the splitting resolution from 0.64 to 0.59 for an isotropic normal in 50 dimensions. Indeed, somewhat surprisingly, raising the sample size leads to a type I error at slightly lower resolution values.

We develop similar formulas for a pair of isotropic normals. In that context, our formulas describe the splitting resolution as a function of distance between the means of the two normals. Putting our results for the single and paired normals together provides a concrete example of an embedding which leads to type I or type II errors. If the means of the paired normals are sufficiently close and the single normal is sufficiently eccentric, then we will not be able to split the paired normals without also splitting the single normal.

Our results have limitations. The form of modularity we consider (1) assumes the configuration model as a null model. While this is the form used in most scRNAseq clustering, other models are possible [24, 20]. The configuration model may not be appropriate for scRNAseq datasets and a different model may lead to different results. We have tested the accuracy of our approximations, however our approximations may not be accurate outside of the parameter regimes we consider. We used a particular scRNAseq clustering workfiow. Other workfiows, for example with a dimension reduction step that is not PCA based [5, 19, 4, 30], may not occupy the same parameter regimes. Similarly, we have only considered knn graphs, while in practice shared nearest neighbor or UMAP graphs are often used. Our assumption that the pair of normals are isotropic limits our ability to analyze scRNAseq dataset since the GMM models we fit are not isotropic. Constructing an approximation for non-isotropic pairs represents a direction of future work.

Overall, our results strongly support the need for hierarchical clustering methods in the context of scRNAseq. Hierarchical modularity clustering has been applied to scRNAseq datasets, especially in the context of large cell atlases, for which practitioners have noted the difficulties of applying modularity clustering across scales [29, 9, 25], but no theory of modularity based hierarchical clustering exists. The development of such a theory represents a direction of future work.

## 3 Methods

### 3.1 Datasets

The count matrix for each dataset was downloaded from public repositories (cellxgene, GEO, and the 10X genomics website). Full download details are available in the Supporting Information. Raw count matrices were filtered by library size, gene count across cells, and fraction of mitochondrial expression. The specific values that we filtered for varied by the dataset and are described in the Supporting Information. For the blood, brain, pancreas and PBMC datasets, we further randomly subsampled to 30, 000 cells for the purposes of reducing computation time. The other datasets had 27, 525 (eye), 17, 786 (heart), and 26, 274 (tongue) cells in the filtered count matrix.

After the dataset specific processing, we applied the following processing pipeline to all datasets. We selected 1000 genes using scanpy’s `highly_variable_genes` function with `flavor=11seurat_v311`, normalized and scaled the data using the residual algorithm in [15], applied a PCA with dimension 50, used sklearn’s `kneighbors_graph` function to form the knn graph, and applied modularity clustering using the Leiden algorithm through the `leidenalg` package. UMAP visualizations were created using scanpy. Python code to reproduce all results in this manuscript is available at,

https://github.com/SLeviyang/resolutionTradeoffs

### 3.2 The Pareto Modularity Frontier

The arguments below apply to any penalized optimization with finitely many solutions. We include them here for the sake of completeness. Let (*g*_*i*_, *h*_*i*_) be optimal *g, h* combinations with the *g*_*i*_ in increasing order. We can assume the *g*_*i*_ are strictly increasing because if there are two optimal clusterings 𝒞, *C*^*′*^ with *g*(𝒞) = *g*(𝒞 ^*′*^) then *h*(𝒞) = *h*(𝒞 ^*′*^) because otherwise one of the clusterings is not optimal. Given a strictly increasing order of the *g*_*i*_, the *h*_*i*_ must be strictly decreasing. Indeed, if the *h*_*i*_ are not in strictly decreasing order than there exists *i* for which *g*_*i*_ *< g*_*i*+1_ and *h*_*i*_ ≤ *h*_*i*+1_, but then *g*_*i*_ + *γh*_*i*_ *< g*_*i*+1_ + *γh*_*i*+1_ for all *γ* and so *g*_*i*+1_, *h*_*i*+1_ cannot be an optimal combination.

For each (*g*_*i*_, *h*_*i*_) combination, let *J*_*i*_ be the set of *γ* values for which *g*_*i*_ +*γh*_*i*_ is optimal. We claim that *J*_*i*_ is a closed interval or a single point. To see this, let *J*_*i*,*l*_ = {*γ* ≥ 0 | *g*_*i*_ + *γh*_*i*_ ≤ *g*_*l*_ + *γh*_*l*_}. The *J*_*i*,*ℓ*_ are of the form [0, *x*] or [*x*, 0] for *x* ∈ *ℝ* ^+^ and 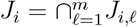. Clearly *J*_*i*_ is either a closed interval, a single point, or the empty set. *J*_*i*_ cannot be the empty set since (*g*_*i*_, *h*_*i*_) is optimal for some *γ*.

If *J*_*i*_ is a single point, then we can remove this (*g*_*i*_, *h*_*i*_) combination since for every *γ* for which (*g*_*i*_, *h*_*i*_) are optimal, then another (*g*_*j*_, *h*_*j*_) is also optimal. (Alternatively, we could have defined the (*g*_*i*_, *h*_*i*_) as the smallest set of optimal combinations for which there is an optimal combination for each *γ* ∈ *ℝ* ^+^.)

At this point we’ve shown that for every (*g*_*i*_, *h*_*i*_) there is a closed interval *J*_*i*_ that contains the *γ* at which (*g*_*i*_, *h*_*i*_) is optimal. We claim that *J*_*i*_ are either disjoint or intersect at a single point.

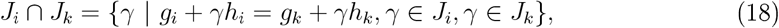

(*g*_*i*_, *h*_*i*_) and (*g*_*k*_, *h*_*k*_) are both optimal for *γ* ∈ *J*_*i*_ ∩ *J*_*k*_. Since *g*_*i*_ + *γh*_*i*_ = *g*_*k*_ + *γh*_*k*_ has a unique solution in *γ, J*_*i*_ ∩ *J*_*k*_ is either empty or contains a single point.

Finally, we claim the *J*_*i*_ are increasing in *i*, meaning that if *γ* ∈ *J*_*i*_, *γ*^*′*^ ∈ *J*_*i*+1_ then *γ* ≤ *γ*^*′*^. To see this, note that if *γ* ∈ *J*_*i*_ then *g*_*i*_ + *γh*_*i*_ ≤ *g*_*i*+1_ + *γh*_*i*+1_, which is solved by *γ* ≤ (*g*_*i*+1_ − *g*_*i*_)*/*(*h*_*i*_ − *h*_*i*+1_). Similarly, if *γ*^*′*^ ∈ *J*_*i*+1_ then noting *h*_*i*_ *> h*_*i*+1_ gives *γ*^*′*^ ≥ (*g*_*i*+1_ − *g*_*i*_)*/*(*h*_*i*_ − *h*_*i*+1_).

### 3.3 Proof of (5)

Let 𝒞 _*b*_ be the baseline cluster, let *n*_*b*_ be the number of nodes in 𝒞 _*b*_, and assume that the baseline cluster is disconnected from the other nodes in the graph.

Consider first the local splitting resolution. Let 𝒞 be a clustering of 𝒞 _*b*_ and let 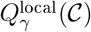 be the modularity restricted to the graph formed by 𝒞 _*b*_. If 𝒞 contains a single cluster, i.e. all the nodes in 𝒞 _*b*_ are put in one cluster, then 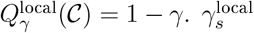 will be the minimum resolution *γ* at which there exists a clustering 𝒞 of 𝒞 _*b*_ containing more than one cluster such that,

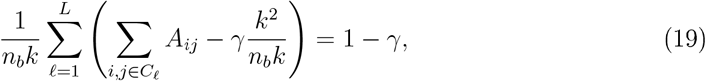

where 𝒞 _1_, 𝒞 _2_, … , 𝒞 _*L*_ cluster 𝒞 _*b*_.

Now consider the global splitting resolution. First note that if 𝒞 is an optimal clustering of the *n* nodes of the graph, then a cluster 𝒞 _*ℓ*_ ∈ 𝒞 contains nodes from a single disconnected component of the graph. Otherwise, we could split 𝒞 _*ℓ*_ according to the disconnected components of the graph and this new clustering would have an equal *g* value and a lower *h* value, contradicting the optimality of 𝒞. We can therefore split optimization of the modularity (1) into an optimization of the modularity restricted to nodes in 𝒞 _*b*_ and to the other nodes in the graph, respectively. Letting 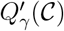 be the modularity restricted to nodes in 𝒞 _*b*_,

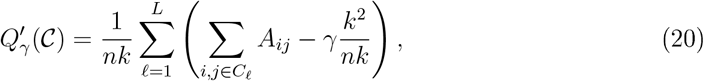

where 𝒞 _1_, 𝒞 _2_, … , 𝒞 _*L*_ cluster 𝒞 _*b*_. If 𝒞 contains a single cluster, 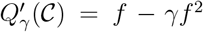, where *f* = *n*_*b*_*/n*. The global splitting resolution *γ*_*s*_ will be the minimum resolution *γ* at which there exists a clustering 𝒞 of 𝒞 _*b*_ containing more than one cluster such that,

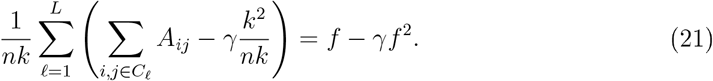

Dividing both sides of the above equation by *f* and using *nf* = *n*_*b*_ gives,

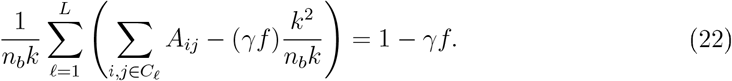

Comparing (19) to (22) gives 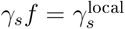.

### 3.4 *γ*_*s*_ for the normal model

Let *x*_*i*_ ∼ 𝒩 (0, Σ) for *i* = 1, 2, … , *n* with each *x*_*i*_ ∈ *R*^*p*^. The restriction to mean zero is not a limitation since we can always shift and rotate the *x*_*i*_ without affecting the distribution of the adjacency matrix *A*. Let *X* be the row binding of the *x*_*i*_ samples, making *X* an *n* × *p* matrix.

To compute *E*[*e*_*i*_] and *E*[*f* ], we need to specify a clustering and we need the distribution of the distances ∥*x*_*i*_ − *x*_*j*_∥^2^ over *j* since these distances determine the k-nearest neighbors of *x*_*i*_. For the clustering, we claim that the projection of the *x*_*i*_ onto the dominant singular vector of *X* gives an approximately optimal clustering. This is the essential idea underlying our results. We do not prove that this clustering is optimal, instead we estimate *γ*_*s*_ under this clustering and the numerical results shown above justify our assumption, at least in the context of the parameter regimes considered.

To make our choice of clustering precise, let 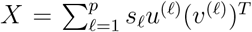 be the svd decom-position of *X* where *s*_*ℓ*_, *u*^(*ℓ*)^, *v*^(*ℓ*)^ are the *ℓ*th singular value and vectors of *X*. (Throughout, for simplicity, we assume *n > p*.) Letting *c*_*i*_ ∈ {1, −1} specify the cluster of *x*_*i*_, we set *c*_*i*_ = sign(*x*_*i*_ *· v*^(1)^) for *i* = 1, 2, 3, … , *n*.

Turning to the distances, a standard linear algebra computation, using the orthogonality of the singular vectors *v*^(*ℓ*)^, gives

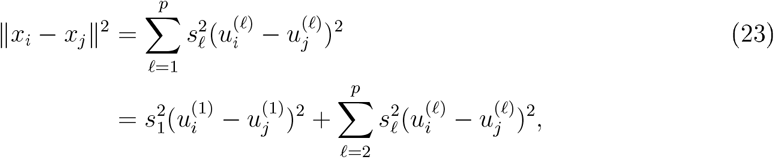

where 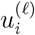 is the *i*th coordinate of *u*^(*ℓ*)^. The expression to the right of the second equality directly above expresses the distance as the sum a clustering and a noise term. Since 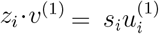, the first summand, 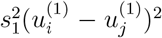, captures all the clustering information, making the second summand a noise term.

Now we apply a sequence of five approximations. The last of these approximation is Approximation 1 above. In introducing the approximations, our goal is to develop a simple probabilistic description of the distances ∥*x*_*i*_ −*x*_*j*_ ∥^2^ over *j* with *i* fixed. As our first approximation, we assume that the left singular vectors *u*^(*ℓ*)^ have independent coordinates distributed as 𝒩 (0, 1*/n*). To motivate this approximation, recall the svd relation *s*_*ℓ*_*u*^(*ℓ*)^ = *Xv*^(*ℓ*)^. Considering a single coordinate of this relation, we have

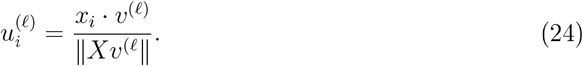

*X* and *v*^(*ℓ*)^ are not independent, but any individual row of *X, x*_*i*_, will be roughly independent of *v*^(*ℓ*)^. Under this approximation, *x*_*i*_ *·v*^(*ℓ*)^ ∼*N* (0, (*v*^(1)^)^*T*^ Σ*v*^(1)^) and *u*^(*ℓ*)^ is a normalization of a vector of iid, normally distributed entries which is well approximated by iid 𝒩 (0, 1*/n*) entries.

As the second approximation, we replace the noise term with a single normal random variable. Recalling that the knn graph is independent up to shifts and rescalings of the *x*_*i*_, let the ∼ symbol represent a distribution equivalent by shifts and rescaling. We can transform the noise term as follows.

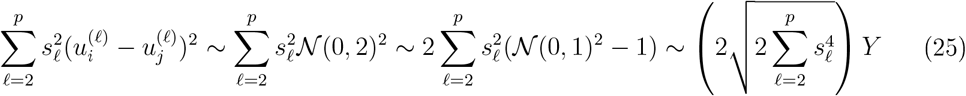

where

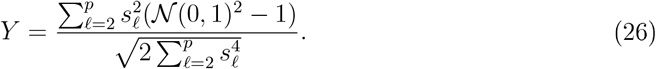

has mean zero and unit variance and is the sum of independent random variables; we can hope that a CLT approximation is accurate and *Y* is approximately a standard normal. We use *Y*∼𝒩 (0, 1) as our second approximation. Applying all these rescalings to the clustering term in (23) as well, we arrive at the following expression for the second approximation,

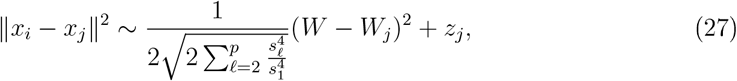

where *W, W*_*j*_, *z*_*j*_ are all 𝒩 (0, 1) and approximate 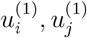 and *Y* respectively. As another rescaling, we multiply the right hand side of (27) by 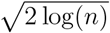, giving

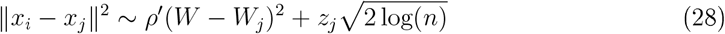

where

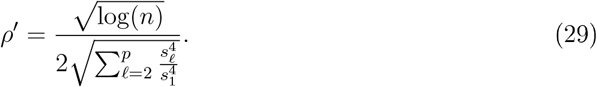

To explain this 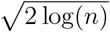 rescaling, we recall a standard order statistic result. Let *z*_*i*_ for *i* = 1, 2, … , *n* be iid samples from a standard normal and let *z*_(*j*)_ be the *j*th order statistic (i.e. the *j*th smallest value of the *n* samples), then for 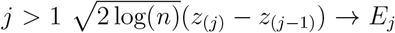 where *E*_*j*_ is an exponential random variable with mean 1*/j*.

The third approximation replaces the 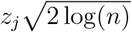 term in (28), by the asymptotic value as *n* → ∞ based on the order statistic analysis. Let *E*_1_, *E*_2_, *E*_3_, … be independent exponential random variables with mean 1, 1*/*2, 1*/*3, … . We approximate *D*_*j*_ ∼ ∥*x*_*i*_ − *x*_*j*_∥^2^ by,

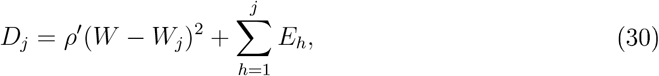

for *j* = 1, 2, 3, … .

The *D*_*j*_ give a scaled and shifted distribution of ∥*x*_*i*_ − *x*_*j*_∥^2^. If we order the *D*_*j*_, the first *k* will correspond to the *k* nearest neighbors on the knn graph Then sign(*W*) and sign(*W*_*j*_) approximate the cluster of *x*_*i*_ and *x*_*j*_, respectively. Putting this together, we finally arrive at an approximation for *E*[*e*_*i*_],

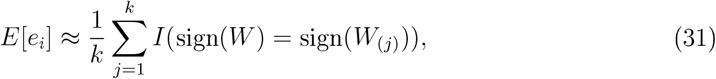

where (*j*) are the order statistics of the *D*_*j*_

The third approximation is essentially Approximation 1 except that *ρ*^*′*^ depends on the singular values of *X*. Our fourth approximation replace the *s*_*i*_ in *ρ*^*′*^ by singular values of a random sample *X*^*′*^ formed from 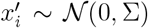. This means that our singular values no longer depend on the data matrix *X*. Finally, as *n*→ ∞ , 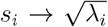 , which gives the *ρ* of Approximation 1.

To estimate *γ*_*s*_, we still need *E*[*f* ], the fraction of *x*_*i*_ with *c*_*i*_ = −1.

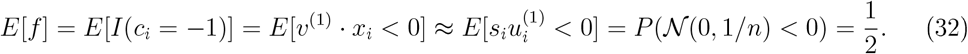

Figure 10 shows the relative accuracy of *γ*_*s*_ estimation after each step in the derivation of Approximation 1. Step 2 causes an upward bias which seems to be corrected by Step 3. There is little difference in accuracy between steps 3, 4 and 5.

**Figure 10.**
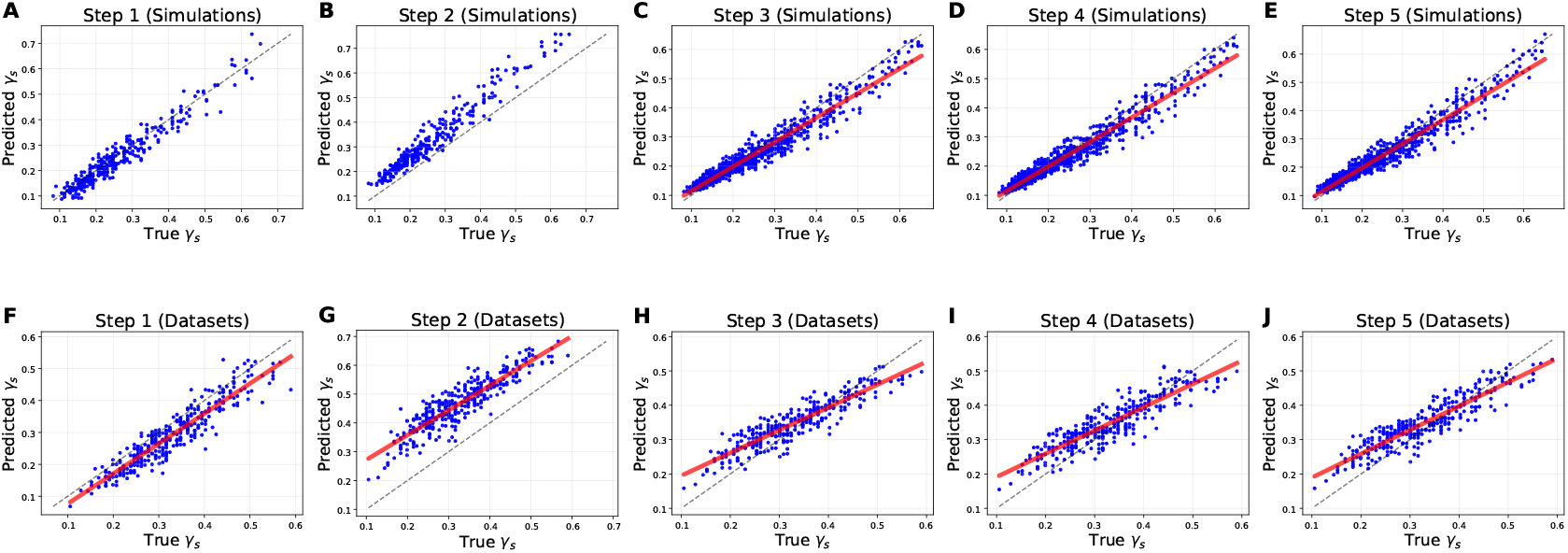
Accuracy over the Sequence of Steps Used to Derive Approximation 1. For each step, the top (bottom) panels show the predicted splitting resolution vs the true splitting resolution for constructed (dataset) generated Σ. We assume that singular vectors are normally distributed (step 1), approximate the noise term by a normal (step 2), rescale and approximate the normal associated with the noise using order statistic asymptotics (step 3), replace the singular values of *X* by the singular values from a resampling (step 4), and replace the singular values by the singular values of Σ (step 5). The approximation of step 5 is Approximation 1

### 3.5 *γ*_*s*_ for the Paired Isotropic Normal Mixture Model

Assume the model,

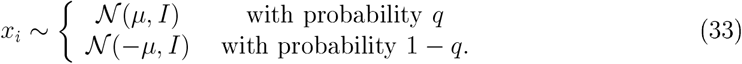

As above, let *X* be the row binding of the *x*_*i*_ for *i* = 1, 2, … , *n. X* has the form,

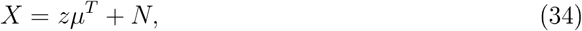

where *z* ∈*ℝ*^*n*^ and *z*_*i*_ = 1 or *z*_*i*_ = −1 depending on whether *x*_*i*_ is drawn from the first or second mixture respectively and *N* is an *n*× *p* matrix with iid standard normal entries. Letting 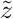 and 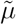 be the normalization of *z* and *µ* to Euclidean length 1,

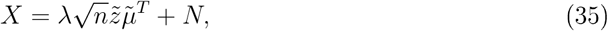

where *λ* = ∥*µ*∥. Then we can consider a rescaled version of *X*,

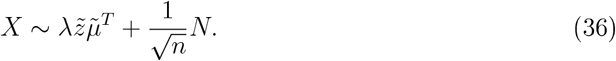

As in the single normal case, we express *X* in terms of its svd and cluster using 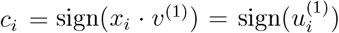. The difference to the single normal case is that we use random matrix theory (RMT) results to characterize the dominant singular value *u*^(1)^. Indeed, (36) is exactly in the form of Johnstone’s spiked model, a well known RMT model [11, 1]. In [2], Benaych-Georges and Nadakuditi derive the distribution of the singular vectors and values for *X* given by (36) in the *n, p→ ∞* limit with *p/n* fixed.

Specifically, as shown in [2],

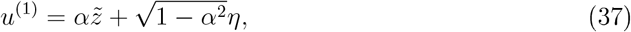

where the entries of *η* are iid 𝒩 (0, 1*/n*), and for *ℓ* = 2, 3, … , *p*, the coordinates of *u*^(*ℓ*)^ are iid 𝒩 (0, 1*/n*). We again go through a sequence of approximations ending in Approximation 2 above. Our first approximation assumes the normality of the *u*^(*ℓ*)^ but the singular values and *α* are computed from the data. Our second and third approximations are essentially identical to the approximations made in the single normal case. We then arrive at Approximation 2, except that *α*^2^ and *ρ* are computed based on a singular value decomposition of *X*.

To remove the parameter dependence on *X*, we make a fourth approximation using results from 2]. The value of *α* is given in equation 11 in [2],

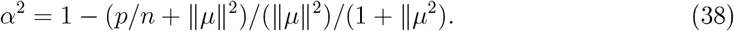

The singular values *s*_*j*_ are given in equation 9 of [2],

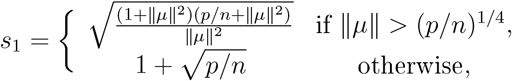

and 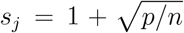 for *j* = 2, 3, … , *p*. Finally, our fifth approximation uses the above formulas for *α*^2^ and *s*_*j*_ with *p/n* = 0.

To find *E*[*f* ], we need to find the fraction of samples in cluster 1. We compute the expected value of this fraction.

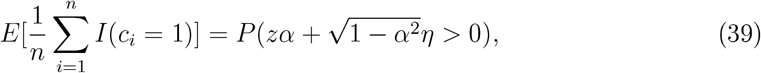

where *z* is Rademacher rv with success probability *q*. A simple calculation gives,

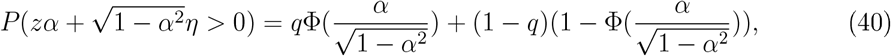

where Φ is the standard normal cdf.

Figure 11 shows the relative accuracy of *γ*_*s*_ estimation after each step in the derivation of Approximation 2. All steps lead to approximations with a downward bias.

**Figure 11.**
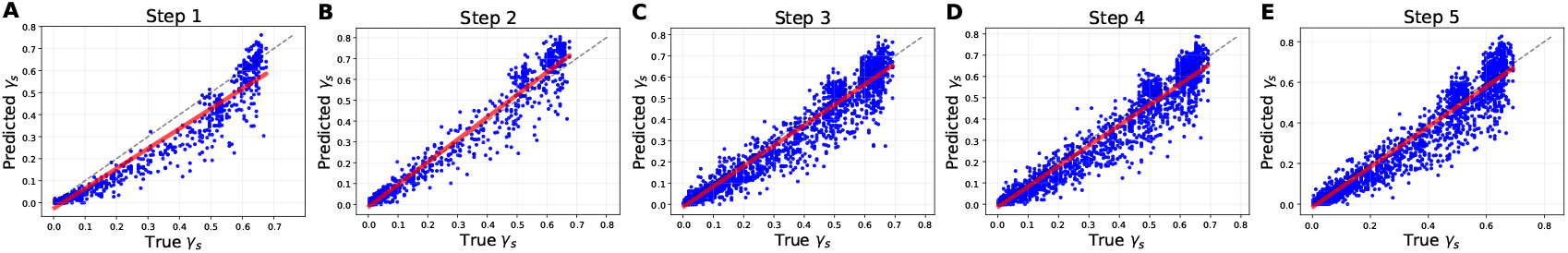
Accuracy over the Sequence of Steps Used to Derive Approximation 2. For each step, the panel shows the estimated splitting resolution vs the true splitting resolution. We assume that singular vectors are normally distributed (step 1), approximate the noise term by a normal (step 2), rescale and approximate the normal associated with the noise using order statistic asymptotics (step 3), replace singular values and *α*^2^ by the formulas of [2] (step 4), and apply *p/n* = 0 to the formulas (step 5). The approximation of step 5 is Approximation 2

**Figure 12.**
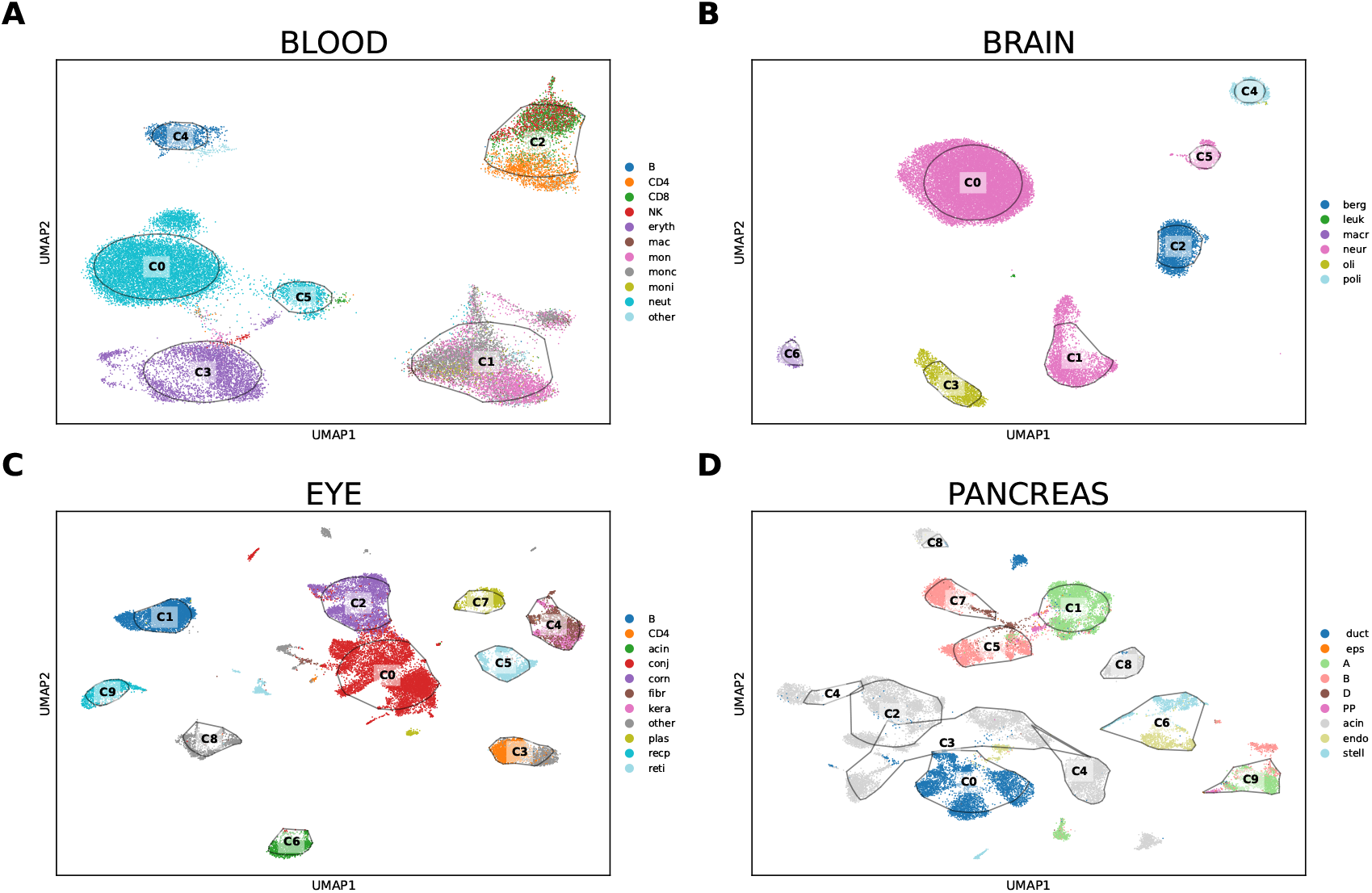
Baseline Clustering. UMAP visualization of baseline clustering for the datasets not shown in Figure 2 of the manuscript.

## Supplementary Information

See the main text for dataset references.

### Brain Dataset

We downloaded the Human Brain Cell Atlas v1.0 - Dissection: Cerebellum (CB) - Cerebellar Vermis - CBV dataset from cellxgene. We filtered for cell counts between 10^3.2^ and 10^4.4^, gene counts between 10^3^, 10^3.8^ and mitochondrial gene expression less than 4%. From these filtered cells, we randomly chose 30, 000 cells.

### Bood Dataset

We downloaded the Tabulas Sapiens - Blood dataset from cellxgene. We filtered for cell counts below 10^4.5^, gene counts between 10^2.3^, 10^3.85^ and mitochondrial gene expression less than 14%. From these filtered cells, we randomly chose 30, 000 cells.

### Eye Dataset

We downloaded the Tabulas Sapiens - Eye dataset from cellxgene. We filtered for cell counts below 10^4.9^, gene counts between 10^2.6^, 10^3.8^ and mitochondrial gene expression less than 17%. This filtering gave 27, 525 cells which we did not subsample.

### Heart Dataset

We downloaded the Tabulas Sapiens - Heart dataset from cellxgene. We filtered for cell counts below 10^4.9^, gene counts between 10^3^, 10^3.8^ and mitochondrial gene expression less than 20%. This filtering gave 17, 786 cells which we did not subsample.

### Pancreas Dataset

We downloaded the pancreas dataset from the Pancreas Atlas website hpap.pmacs.upenn.edu. The dataset has accession GSE148073. We filtered for cell counts between 10^3.2^ and 10^5^, gene counts between 10^2.6^, 10^3.8^ and mitochondrial gene expression less than 10%. From these filtered cells, we randomly chose 30, 000 cells.

### PBMC Dataset

We downloaded the PBMC dataset from the 10x genomics website, http://support.10xgenomics.com/single-cell/datasets

We downloaded the annotations for the dataset from, https://github.com/10XGenomics/single-cell-3prime-paper/tree/master

We filtered for cell counts between 10^3^ and 10^3.5^, gene counts greater than 50 and mitochondrial gene expression less than 5%. We subsampled 30, 000 cells.

### Tongue Dataset

We downloaded the Tabulas Sapiens - Tongue dataset from cellxgene. We filtered for cell counts below 10^4.9^, gene counts between 10^2.9^, 10^4^ and mitochondrial gene expression less than 17%. This filtering gave 26, 274 cells which we did not subsample.

